# Molecular-level analysis of serum IgG repertoires in COVID-19-vaccinated people with cystic fibrosis identifies abundant convergent antibodies

**DOI:** 10.64898/2026.07.08.737356

**Authors:** Steven Ionov, Ruth I. Connor, Nicholas C. Curtis, Seungmin Shin, Brian H. Kim, Niklas Koehne, Ji-Ye Park, Alix Ashare, Peter F. Wright, Jiwon Lee

## Abstract

**Background:** People with cystic fibrosis (pwCF) are at increased risk of severe disease following respiratory viral infections including influenza and respiratory syncytial virus, and were therefore given priority for COVID-19 vaccination. Retrospective epidemiological data have revealed a lower incidence of SARS-CoV-2 infection and a reduced fatality rate among pwCF than in the general population, possibly from the stringent infection control measures and early adoption of protective behaviors. Despite several reports of adequate binding and neutralizing titers after vaccination, the molecular features of vaccine-elicited serum antibody repertoires in pwCF remain unknown.

**Methods:** We performed high-resolution proteomic analysis of serum IgG, combined with next-generation sequencing of B cells, to quantitatively profile Spike (S)-reactive serological repertoires from nine infection-naïve pwCF after two doses of COVID-19 mRNA vaccine. We recombinantly expressed 20 IgG clonotypes from 6 pwCF as monoclonal antibodies (mAbs) and measured their binding and neutralization against vaccine-strain and variant viruses.

**Results:** All donors mounted strong binding and neutralizing titers to vaccine-strain virus. Ig-Seq revealed diverse serum repertoires in all vaccinees, with antibodies targeting the receptor-binding domain (RBD) comprising 64% of each circulating repertoire by abundance. Several serum IgG clonotypes identified across our cohort shared identical or similar *IGHV* and CDRH3 amino acid sequences with serum IgG from other pwCF or S-reactive mAbs isolated from non-CF individuals. These ‘convergent’ antibodies made up 14.6% of the anti-S repertoires in our cohort, reaching 24.8% in one pwCF. Convergent antibodies tended to be RBD-reactive and enriched in specific *IGHV*; surprisingly, a higher abundance of convergent antibodies in serum correlated with a lower breadth of serum neutralization. All 20 mAbs bound Wuhan S with high affinity (EC_50_ < 10 nM), and only RBD-reactive mAbs conferred neutralization to vaccine or variant pseudovirus. While most mAbs bound both the B.1.617.2 (Delta, 17/20) and B.1.1.529 (Omicron, 12/20) strains, convergent mAbs were less likely to bind either variant.

**Conclusions:** Post-vaccine serum IgG repertoires in pwCF are dominated by RBD-focused, high-affinity antibodies and include a substantial convergent component shared with non-CF vaccinees. These findings demonstrate that pwCF mount antibody responses comparable to the general population, and a large group of convergent antibodies may contribute to strain-specific rather than cross-variant immunity.

## Introduction

Cystic fibrosis (CF) is a life-limiting genetic disorder caused by mutations in the Cystic Fibrosis Transmembrane Conductance Regulator (*CFTR*) gene[1]. CF pathology involves progressive deterioration of lung function and periodic pulmonary exacerbations, which are associated with worsening disease and facilitate colonization with CF pathogens such as *Pseudomonas aeruginosa*[2–7]. Respiratory viral infections are responsible for over 40% of pulmonary exacerbations in people with CF (pwCF)[8], highlighting the importance of immune protection from viruses in this population. pwCF were considered to be at increased risk when SARS-CoV-2 emerged[9] and were recommended priority access to COVID-19 vaccines[10–12]. Surprisingly, incidence of SARS-CoV-2 infection among pwCF was lower than among non-CF individuals, and fatality rates among non-transplanted pwCF in the pre-vaccine period were over twofold lower among pwCF than the general population (0.7% vs. 1.7%, respectively)[13–15]. This finding was especially unexpected in light of the higher disease severity and mortality rates among pwCF after influenza infection[2, 3, 5, 7, 8, 16, 17]. pwCF are known to adhere more strictly to infection-preventive behaviors than the general population, which may partially explain their lower incidence of infection[18]. Nevertheless, the robustness of humoral immune responses in pwCF remains uncertain, with historical and recent studies reporting altered titers of total IgG and differences in B cell populations[19–22].

In the general population, the threat to public health posed by SARS-CoV-2 led to extensive efforts to characterize antibody responses to vaccination. Two-dose mRNA vaccination against ancestral (Wuhan) SARS-CoV-2 consistently elicited strong IgG binding titers against the Spike (S) protein and strong neutralizing titers[23, 24]; both of these were later understood to be correlates of protection against infection and partially explain high vaccine efficacy[23–25]. In addition, serology studies showed that the receptor-binding domain (RBD) is an immunodominant region of S[25, 26], and a recent study suggests that the majority of anti-S IgG in non-CF vaccinees is RBD-reactive[27, 28].

Studies of convalescent and vaccinated donors have found groups of S-reactive antibodies with similar complementarity determining region (CDRH3) amino acid sequences[29–33]. These ‘convergent’ antibody groups also feature equal or similar (*IGHV*) genes[34, 35] and tended to bind to the same regions of S. Well known examples include antibodies encoded by *IGHV3-53/3-66* or *IGHV1-24* genes, which target RBD and N-terminal domain (NTD), respectively; both neutralize Wuhan strain, leading to early speculation that convergent responses are protective[36–38]. Further study showed that some convergent antibodies were non-neutralizing to Wuhan or lost neutralization to viral variants of concern (VOCs)[29], [34, 39, 40]. In addition, while convergent antibodies have been identified from B cell sequencing studies, their contribution to the circulating antibody response has not been quantified, in part because not all circulating antigen-specific B cells secrete antibody into serum[41], precluding an understanding of whether they enable or inhibit robust neutralizing responses.

Despite extensive characterization of COVID-19 vaccine antibody repertoires in the general population, similar studies in pwCF have largely been limited to measurements of post-vaccine antibody binding titers[42–45]. The molecular features of S-reactive circulating repertoires in CF are unknown, and whether pwCF elicit antibodies with similar sequences as non-CF vaccinees is not understood. This presents a need to investigate key features of circulating repertoires in pwCF, including the identities of circulating IgG, their preferred targeting regions, and the binding and neutralizing properties of the individual antibodies that comprise them. Resolving these features in the poorly studied context of pwCF would clarify how antibody responses contribute to clinical outcomes after COVID-19, and would show where those responses resemble or differ from the general population.

To address these gaps, we used Ig-Seq[46] to profile the circulating IgG repertoires to S in 9 pwCF after two doses of Wuhan mRNA vaccine. RBD-reactive antibodies dominate the anti-S repertoire, and 60% of the unique antibodies targeted RBD. Repertoire convergence was present across all 9 donors, and most convergent antibodies were also similar to anti-S mAbs isolated from non-CF individuals. Convergent antibodies comprised up to 24.8% of circulating anti-S IgG antibodies, and their abundance correlated inversely with the breadth of neutralization to VOCs. By expressing 20 monoclonal antibodies (mAbs) from across 6 pwCF, we found that all bound Wuhan S with high affinity; convergent antibodies were less likely to bind to both Delta and Omicron S. pwCF thus elicit robust responses that resemble those of non-CF vaccinees at both the repertoire and single-antibody levels. By quantifying convergent antibodies directly in serum, we further show that they make up a substantial share of the circulating response and that their abundance tracks with reduced neutralization breadth, a relationship relevant to vaccine responses in CF and the general population alike.

## Results

### pwCF mount robust binding and neutralization titers upon COVID-19 vaccination

Serum samples were collected from nine infection-naïve pwCF, between the ages of 19 and 60, at 28-31 days after their second dose of either the BNT162b2 or mRNA-1273 vaccine (**Fig. 1a**, **Table S1**). Total serum IgG binding titer against the Wuhan S were detectable in all donors, with half-maximal serum binding dilution (EC_50_) ranging from 1.2 × 10^3^ to 4.3 × 10^3^ (geometric mean 2.0 × 10^3^; **Fig. 1b**). Serum 50% neutralizing titers (NT_50_), measured using a vesicular stomatitis virus pseudotyped with Wuhan S (VSV-Wuhan), ranged from 497 to 2747 (geometric mean NT_50_, 907; **Fig. 1c**), which is comparable to those reported in vaccinated non-CF individuals (approximately 500-1100)[47–50]. Higher S-reactive IgG EC_50_ correlated with higher serum neutralizing titers (Spearman ρ = 0.783, *p* = 0.017; **Fig. 1d**), consistent with observations in non-CF cohorts[51]. Additionally, neither binding nor neutralization titers showed any age-related differences in this small cohort (**Fig. S1**).

**Figure 1.**
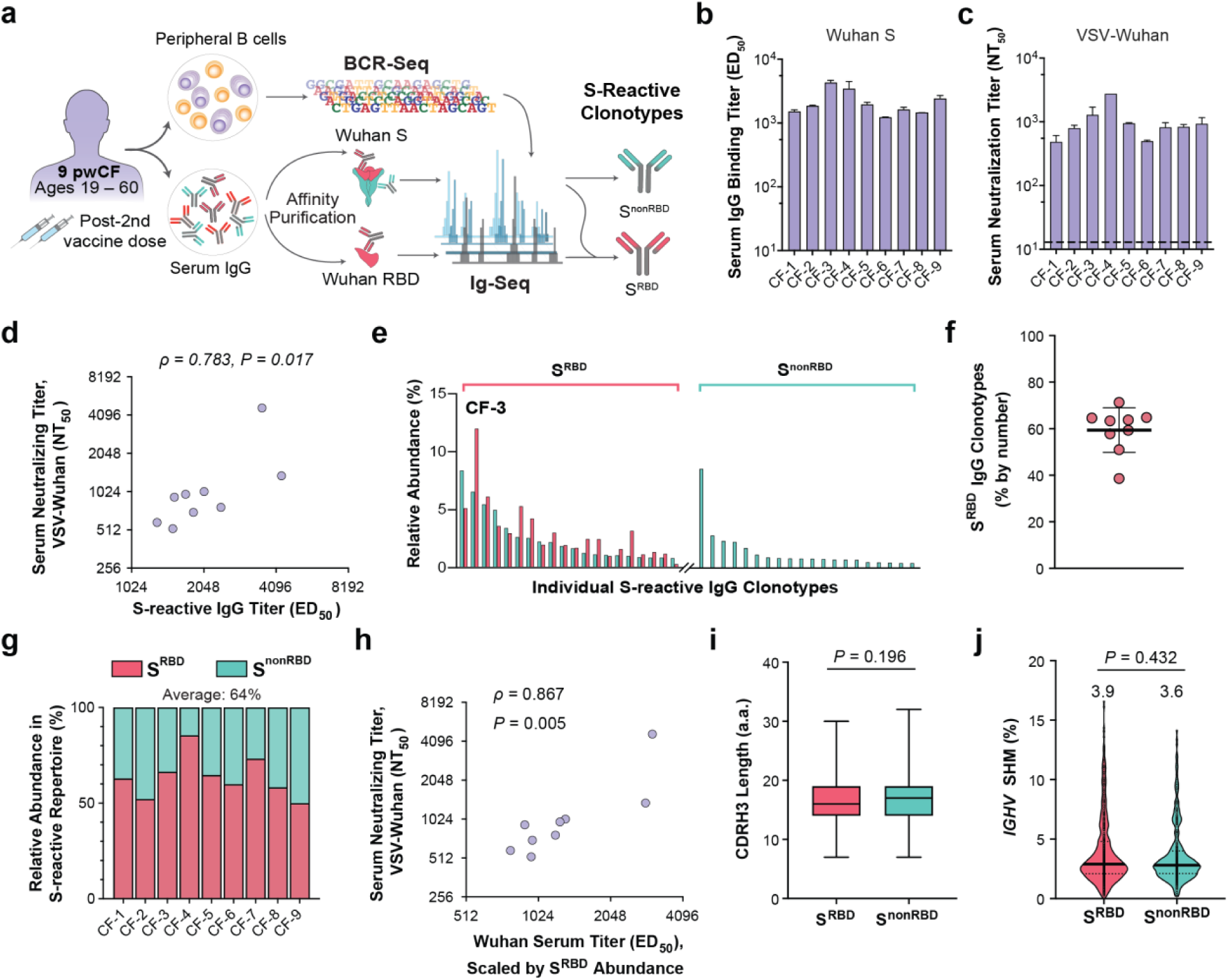
RBD-reactive antibodies dominate post-vaccine serum repertoires in pwCF. **a.** Study design. Samples collected from nine pwCF were processed via BCR-Seq and Ig-Seq. S-reactive IgG clonotypes were classified as S^RBD^ and S^nonRBD^, based on their presence and absence in the RBD dataset, respectively. **b.** Total serum IgG binding titers to S, measured by ELISA and expressed as half-maximal binding dilutions (ED_50_). **c.** Total serum 50% neutralization titers (NT_50_) against VSV-Wuhan. Baseline (12.5) represents lowest dilution used for measurement of NT_50_. **d.** Spearman correlation between total serum neutralization titers and IgG binding titers to S. **e.** Representative histogram showing the top 20 S^RBD^ and S^nonRBD^ clonotypes by relative abundance in donor CF-3. Each bar represents abundance of an individual clonotype; green bars denote abundance of individual clonotypes in the S pulldown, while red bars denote abundances in the RBD pulldown. **f.** Fraction of clonotypes identified as S^RBD^ among all S-reactive unique IgG clonotypes detected in each serum repertoire, quantified by number. **g.** Relative abundances of S^RBD^ IgG in the circulating serum repertoire of each donor. Average is shown above the plot. **h.** Spearman correlation between S-reactive IgG binding titers, scaled by S^RBD^ clonotype abundance, and serum NT_50_ against VSV-Wuhan. **i.** CDRH3 length comparison between S^RBD^ and S^nonRBD^ clonotypes. Box represents first and third quartiles around the median (center line); whiskers range from minimum to maximum. **j.** *IGHV* somatic hypermutation (SHM) levels of serum clonotypes. All clonotypes are divided among S^RBD^ and S^nonRBD^. Solid and dotted lines represent medians and quartiles, respectively. Mean values are displayed. For **b**, **c**, and **f**, error bars indicate SD. For **i** and **j**, statistical significance was measured by Mann-Whitney test.

### RBD-reactive antibodies dominate post-vaccine serum antibody responses

To characterize individual IgG clonotypes within the S-specific antibody repertoires, we performed Ig-Seq[46] on each donor’s serum IgG (**Fig. 1a**). From each serum sample, S-reactive serum IgG was isolated via affinity purification, which were analyzed by high-resolution liquid chromatography-tandem mass spectrometry (LC-MS/MS). The resulting MS spectra were annotated using a matched, donor-specific proteomic search database built from B cell receptor (BCR) sequences from peripheral B cells collected 7 days after second vaccine dose. Because most neutralizing antibodies in the general population target RBD, we performed parallel IgG affinity purifications on all pwCF IgG using the RBD and SD1 subdomain (RBD-SD1, “RBD” henceforth). As illustrated by donor CF-3, who exhibited the highest anti-S serum binding titer, clonotypes were classified as “S^RBD^” or “S^nonRBD^” based on their presence in the RBD purification eluates (**Fig. 1e**). Across all donors, S-specific serum repertoires were diverse, comprising 53-179 unique clonotypes each, and were encoded by a wide range of heavy chain variable region (*IGHV*) genes (**Fig. S2**). Several frequently used genes, including IGHV3-30, IGHV5-51, and IGHV1-69, match those commonly encoding S-reactive antibodies in the general population. This clonotypic diversity is in line with S-reactive repertoire diversities in non-CF donors infected with or vaccinated against SARS-CoV-2[27, 28] and influenza[41, 52]; repertoires were also dominated by a few highly abundant clonotypes, as has been observed in non-CF individuals.

Across all pwCF, S^RBD^ IgG clonotypes accounted for, on average, 60% of S-specific IgG clonotypes by number (range: 39%-71%; **Fig. 1f**). Given that RBD comprises only about 220 of the 1,273 residues of Wuhan S, this concentration of clonotypes marks RBD as the immunodominant region of the vaccine antigen, as described in non-CF individuals. By abundance, S^RBD^ clonotypes constituted an average of 64% of circulating anti-S IgG (range: 50-85%; **Fig. 1g**). Moreover, when we scaled anti-S serum binding titers by S^RBD^ abundance, the correlation with neutralization activity was strong (Spearman ρ = 0.867, *p* = 0.005; **Fig. 1h**).

We did not find any strong preference for a single *IGHV* gene by S^RBD^ clonotypes, though this group showed a mild trend toward preferential *IGHV3-53* and *IGHV3-9* usage (**Fig. S3**), consistent with well-described RBD-binding monoclonal antibodies (mAbs) identified in non-CF populations[53, 54]. In addition, CDRH3 amino acid lengths were comparable between S^RBD^ and S^nonRBD^ groups (16.8 vs. 17.1; **Fig. 1i**), and somatic hypermutation (SHM) levels were likewise similar (3.9% vs 3.6%; **Fig. 1j**). The low overall SHM (**Fig. S4**) implies that circulating anti-S antibodies are newly elicited by naïve B cells[47] rather than reactivated memory B cells. Overall, pwCF mounted robust S-reactive antibody responses that closely resembled those of non-CF populations, with RBD as the dominant target, and showed no clear restrictions in *IGHV* gene usage or CDRH3 features.

### Convergent serum IgG responses are observed across pwCF

Numerous studies have reported convergent B cell clones, defined by similar *IGHV/J* gene usage and CDR3 sequences, across multiple individuals in the general population following vaccination and infection[29, 35, 37, 39, 54–56]. However, convergent responses at the level of circulating IgG clonotypes have not been extensively characterized. We therefore investigated whether pwCF exhibit convergent IgG responses in their serum repertoires.

To do so, we first grouped all 929 IgG clonotypes observed in our cohort by identical *IGHV* gene family, *IGHJ* gene, and CDRH3 length. Within each group, CDRH3 sequences were clustered using an 80% amino acid identity cutoff, and we defined clonotypes clustered between two or more individuals as “convergent IgG clonotypes”. Overall, we identified 16 such groups across the nine pwCF (**Table 1**). Clonotypes within each group shared the same regional binding specificity (S^RBD^ or S^nonRBD^). One group was shared across five of the nine donors and two groups across three donors, while the remaining groups were each shared between two donors. Convergent IgG clonotypes were detectable in circulation of every donor (**Fig. 2a**), and in some instances, including donor CF-4, the most abundant IgG clonotypes were classified as convergent. Across pwCF, these convergent IgG clonotypes constituted an average of 8.2% of the circulating S-specific repertoire by abundance (range: 1.6% – 17.8%; **Table S2**). The substantial sharing of clonotypes even within this small cohort motivated us to ask how far this convergence extends to the broader, predominantly non-CF antibody response.

**Figure 2.**
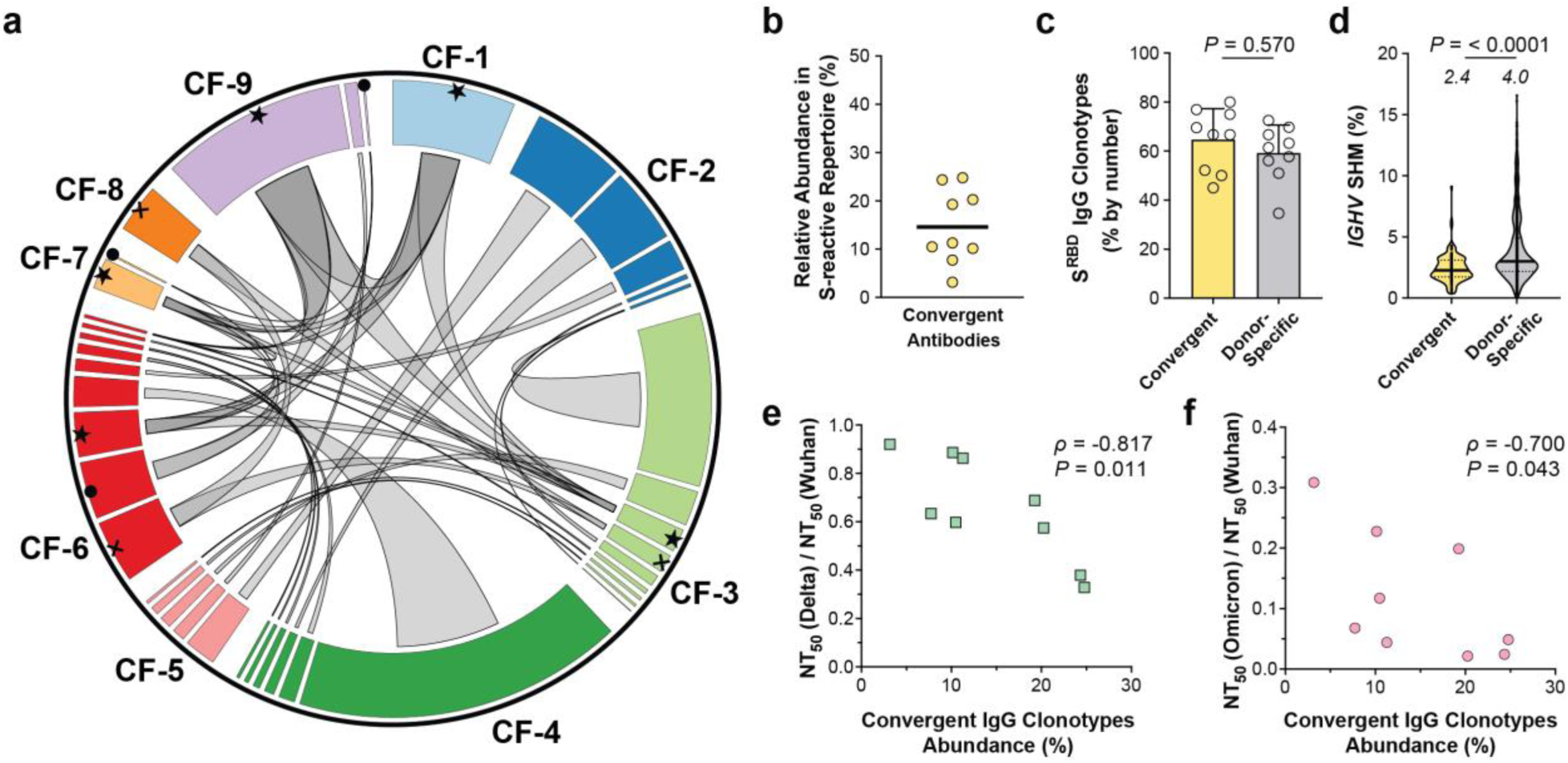
Convergent IgG clonotypes are observed across all pwCF and abundant in serum. **a.** Circos plot of convergent IgG clonotypes across multiple pwCF. Clonotypes belonging to convergent IgG clonotypes are labelled as sectors of different colors; each sector represents a single clonotype and connections are drawn between clonotypes belonging to the same convergent IgG clonotypes. Sector widths are scaled by the relative serum abundance of the corresponding clonotypes. **b.** Relative abundance of convergent IgG clonotypes, including those which were convergent across pwCF and/or with non-CF antibodies (hCoV-AbDab). Horizontal line indicates mean. **c.** Fraction of convergent IgG clonotypes classified as S^RBD^, quantified by number of clonotypes. Each dot represents the fraction in a single donor, and error bars represent SD, respectively. **d.** *IGHV* SHM levels of convergent and donor-specific serum IgG clonotypes. Each serum clonotype in each group is pictured as an individual circle. Solid and dotted lines represent medians and quartiles, respectively. Mean values are listed above each plot. Statistical significance was calculated using Mann-Whitney U-test. **e-f.** Spearman correlations between convergent IgG clonotype abundance and neutralization breadth indices (NBI) for Delta (**e**) and Omicron (**f**), defined as (NT_50_, variant) / (NT_50_, Wuhan).

**Table 1.**
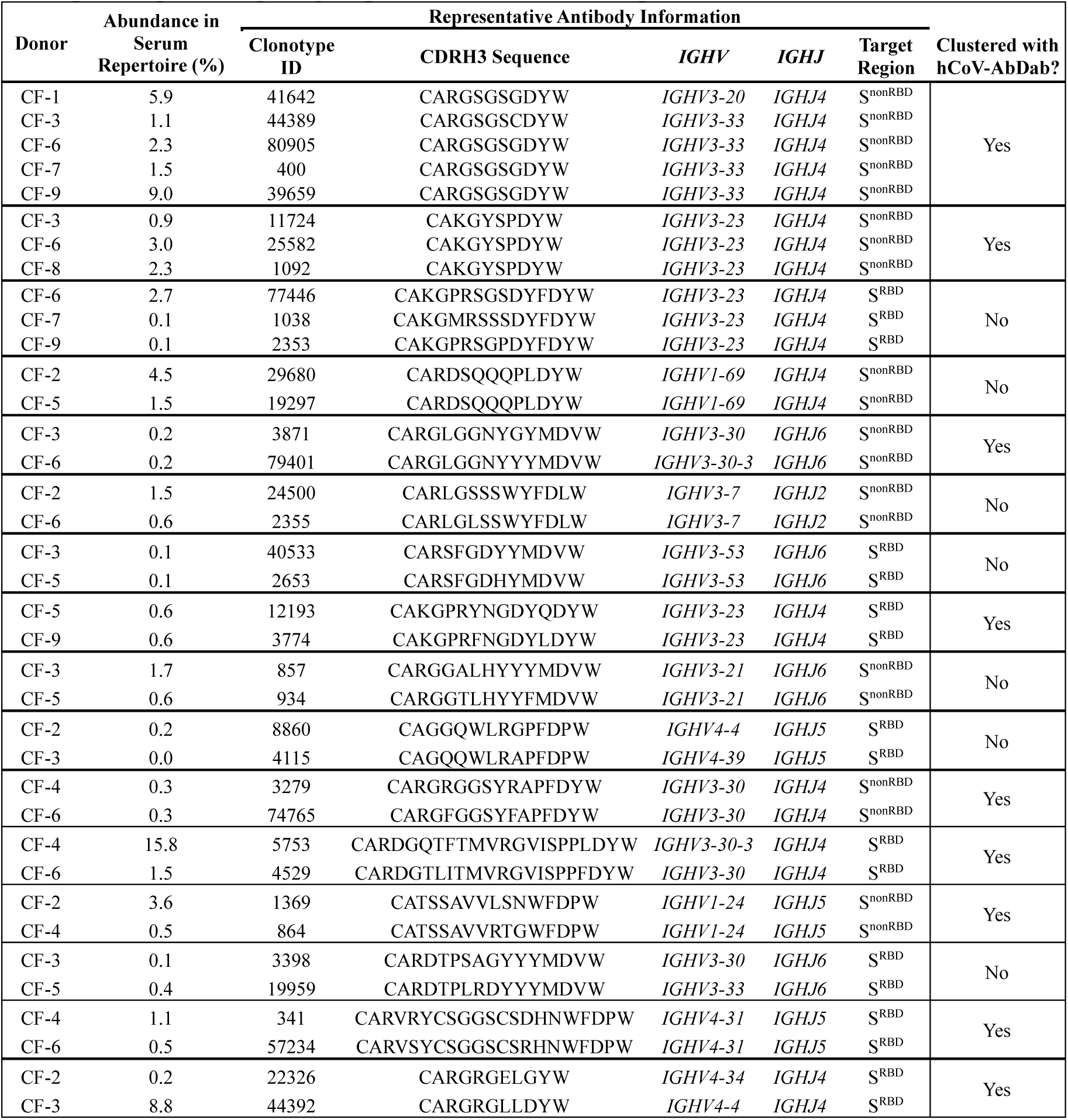
Convergent IgG clonotypes appearing in serum repertoires of multiple pwCF. Clonotype IDs, CDRH3 sequence, *IGHV/IGHJ* usage, and target region are determined from Ig-Seq data. Abundance value indicates total relative abundance of all serum IgG included in the corresponding convergent group, within the donor’s repertoire.

To compare convergent responses in pwCF with those of non-CF individuals, we performed the same grouping and clustering with all 929 clonotypes, along with an additional set of 9,411 S-specific monoclonal antibodies (mAbs) from the human Coronavirus Antibody Database (hCoV-AbDab)[57]. This expanded analysis yielded a total of 81 groups of convergent IgG clonotypes encompassing 127 individual clonotypes. We noted that 9 of the original 16 groups of convergent IgG clonotypes found across multiple pwCF were also clustered with mAbs in the hCoV-AbDab, suggesting that the convergent IgG responses identified in pwCF overlap with those of the general population. Under this broader definition, which counts clonotypes shared across pwCF and/or with general-population antibodies, convergent IgG clonotypes contributed on average 14.6% (range: 3.2% – 24.8%) of each donor’s serum IgG repertoire by abundance, up from 8.2% when counting only clonotypes shared across pwCF (**Fig. 2b**); in donors CF-2 and CF-4, they comprised 24.3% and 24.8% of the repertoire, respectively, underscoring their potential to substantially impact the anti-S serum response.

### Convergent serum IgG responses correlate with lower serum neutralizing breadth

Next, we examined the general features of the convergent IgG clonotypes identified in pwCF. Approximately 65% of these convergent clonotypes were classified as S^RBD^ (**Fig. 2c**), similar to the overall proportion of S^RBD^ clonotypes (**Fig. 1f**). In comparison to non-convergent (“donor-specific”) clonotypes, convergent clonotypes had shorter CDRH3 (**Fig. S5**) and significantly lower *IGHV* SHM levels (**Fig. 2d**). They were also enriched for *IGHV3-23*, *IGHV3-30*, and *IGHV3-53* (**Fig. S6**). Taken together, these findings suggest that convergent IgG clonotypes arise from germline-encoded structural features rather than extensive affinity maturation, thereby explaining how similar antibodies can develop in multiple individuals[54].

To assess the functional impact of these convergent IgG clonotypes against SARS-CoV-2, we performed serum neutralization assays against VSV pseudotyped with Delta and Omicron variants. All nine pwCF had detectable neutralization titers against Delta, and eight of nine (89%) against Omicron (**Fig. S7**). To measure neutralizing breadth, we defined a neutralizing breadth index (NBI)[58–60] against each variant for each donor, by dividing the serum 50% neutralizing titer (NT_50_) to each variant by the NT_50_ to Wuhan.

Although the proportion of convergent IgG clonotypes did not correlate with the magnitude of NT_50_ titers to any given strain, donors with a greater circulating abundance of convergent IgG clonotypes had lower NBI to Delta (Spearman ρ = −0.817, *p* = 0.011) and to Omicron (Spearman ρ = −0.700, *p* = 0.043, **Fig. 2e,f**). In other words, a higher abundance of convergent IgG clonotypes was associated with reduced serum neutralizing breadth, but not with the magnitude of neutralization against any single strain. This pattern is consistent with convergent antibodies binding Wuhan S strongly while remaining sensitive to variant mutations, such that a repertoire weighted toward them neutralizes the ancestral strain well yet loses breadth against VOCs.

### Convergent mAbs exhibit similar binding potency, but may be less VOC-reactive

To examine post-vaccine serum repertoires in pwCF at the level of individual mAb, we expressed 20 representative mAbs recombinantly from the repertoires of 6 donors (**Table 2**). These mAbs represented some of the most abundant IgG clonotypes in serum, such as CF9-1 (22.5% of donor CF-9’s repertoire), and 9 of the 20 mAbs were selected from the convergent IgG clonotypes. These convergent mAbs were drawn both from clonotypes shared across pwCF (**Table 1**) and from clonotypes that clustered with general-population antibodies in the hCoV-AbDab. All 20 mAbs bound Wuhan S with low- to sub-nanomolar affinity, with EC_50_’s as low as 0.04 nM (**Fig. 3a**). Notably, all mAbs derived from S^RBD^ clonotypes (14/20) bound to the RBD construct used for Ig-Seq, which contained both RBD and the small SD1 domain immediately C-terminal to the RBD. Two S^RBD^ mAbs (CF2-56 and CF5-2) bound to this construct but not RBD alone, indicating that these mAbs most likely target SD1 (**Fig. S8**).

**Figure 3.**
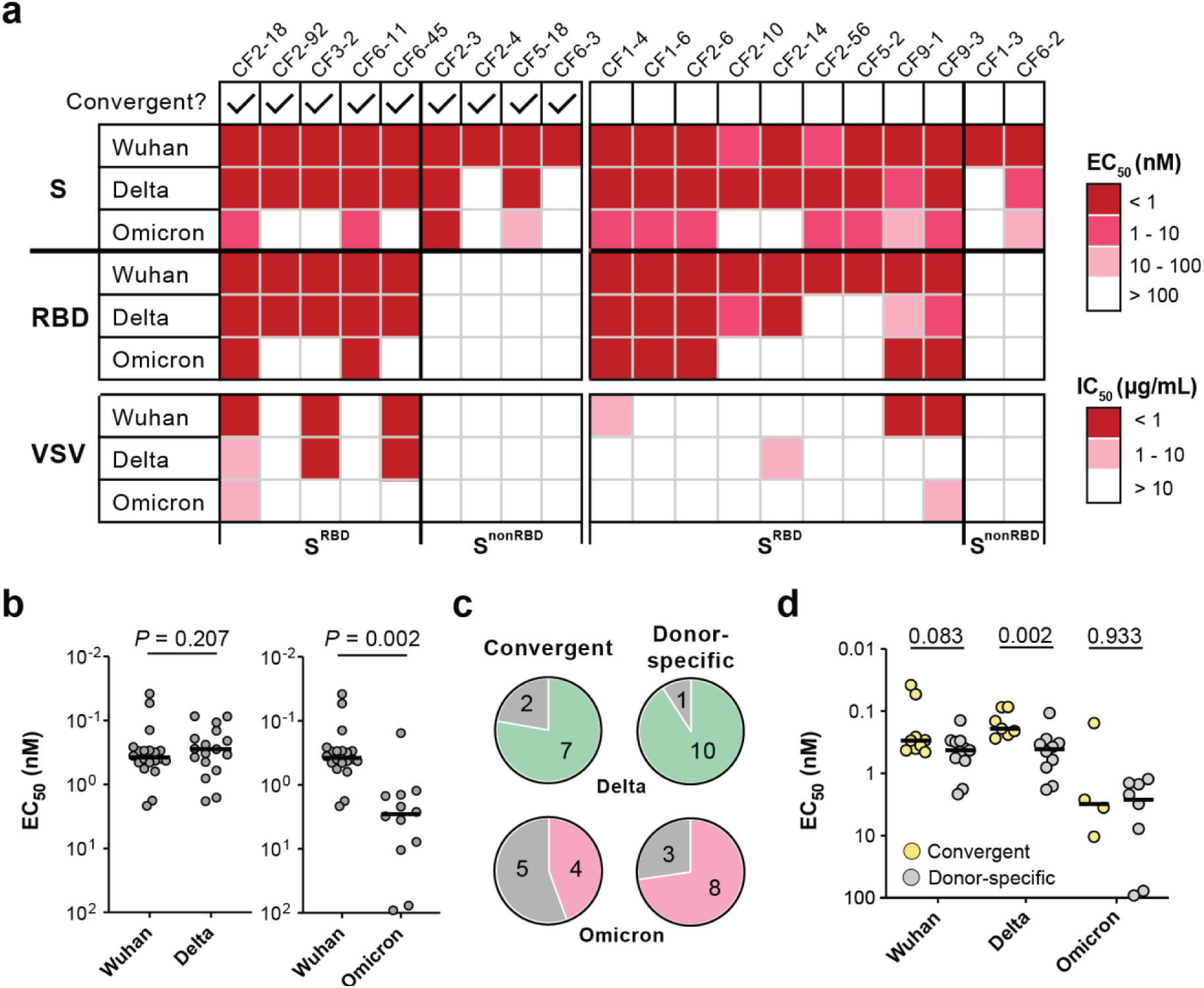
Convergent and donor-specific mAbs bind Wuhan S potently, but convergent mAbs show reduced cross-reactivity to VOC. **a.** Binding and neutralizing properties of each mAb. Top: half-maximal binding concentration (EC_50_) values of all mAbs against Wuhan and VOC S and RBD antigens as determined by ELISA. Bottom: half-maximal inhibitory concentrations (IC_50_) of each mAb against Wuhan and VOC. Color scale indicates binding or neutralizing potency. **b.** Among mAbs that were VOC-reactive (binding EC_50_ of < 100 nM to each VOC), comparison of EC_50_ values to Wuhan and Delta (left) and Omicron (right) S. Statistical significance was measured by Wilcoxon signed-rank tests. **c.** Fraction of all convergent (9) and donor-specific (11) mAbs which bind to VOC S antigens. Pie charts represent fraction of total. Number of mAbs corresponding to each section is indicated. **d.** EC_50_ values of all mAbs to Wuhan and VOCs, split by convergent (yellow) and donor-specific (gray).

**Table 2.**
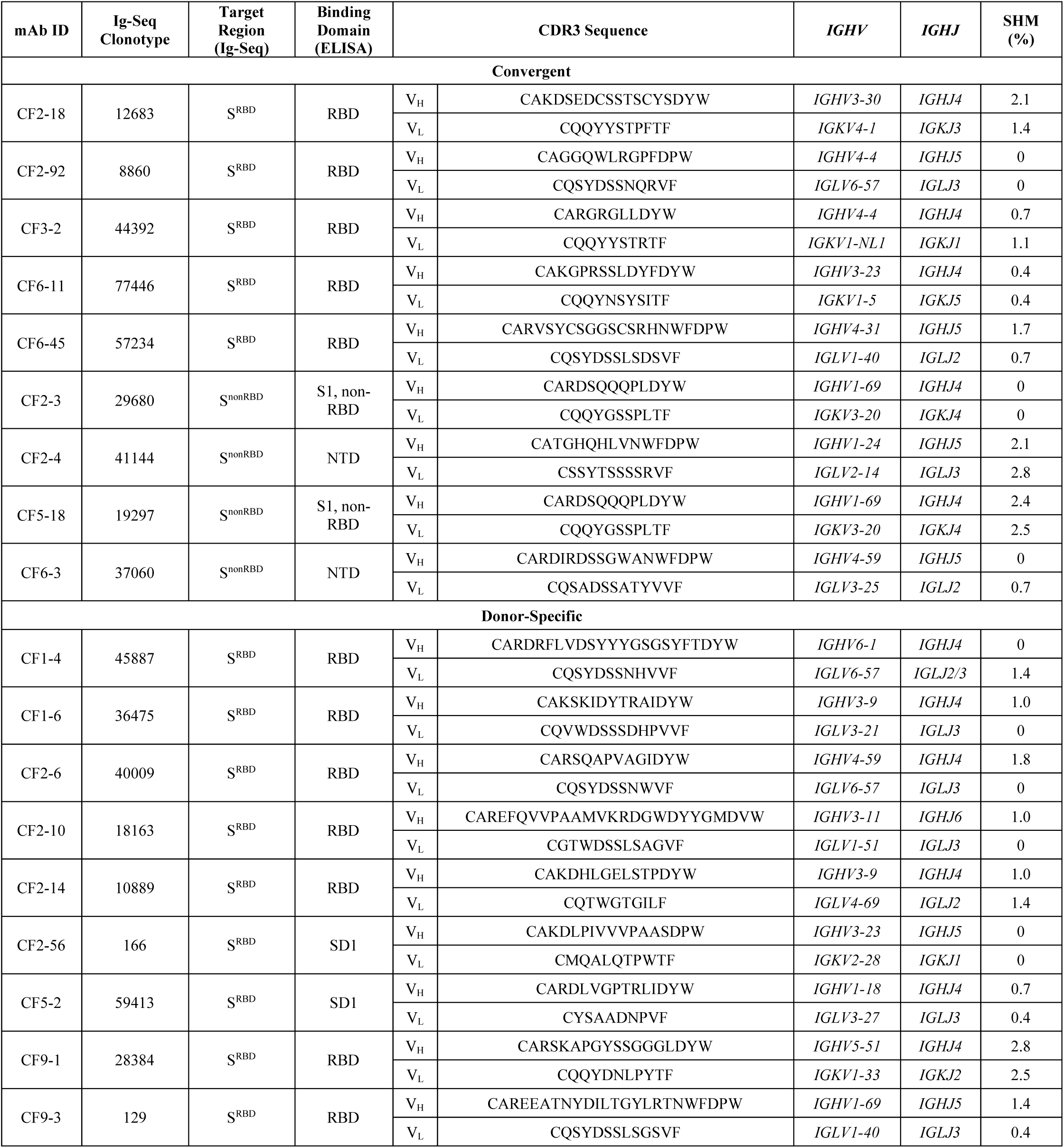
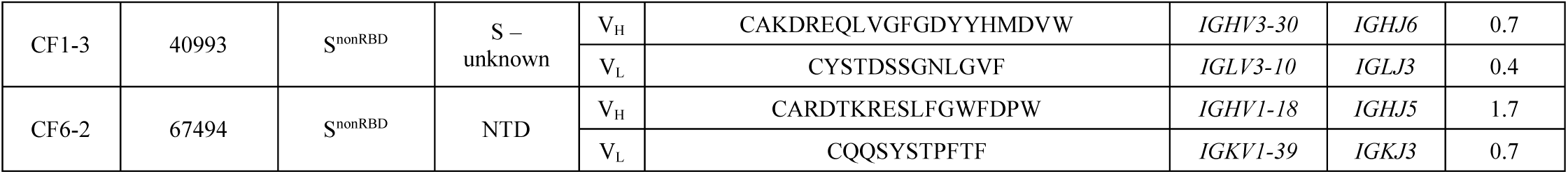
All mAbs expressed for functional assays. mAb ID represents the rank (by abundance) of the corresponding clonotype within each repertoire (*e.g.*, “CF1-4” denotes the clonotype with the fourth highest relative abundance in the repertoire of CF-1).

We next characterized the antigenic regions targeted by these mAbs in more detail. Overall, 19/20 mAbs (95%) bound to regions of the S1 subunit, but targeted diverse regions; apart from the 14 S^RBD^ mAbs, three S^nonRBD^ mAbs bound NTD and two bound to a region of S1 outside of RBD or NTD (**Table S3**). mAb CF1-3 bound full-length S but neither the S1 nor S2 subunits alone, indicating that it targets an epitope present only on full-length S. Our results indicate that S^nonRBD^ mAbs engage diverse regions of S1. To assess binding breadth, we tested each mAb against Delta S and Omicron S (**Fig. 3a**). 17 of 20 (85%) bound Delta S, and 12 of 20 (60%) bound Omicron S. While Delta-binding mAbs generally mirrored the EC_50_ values observed with Wuhan S, Omicron-reactive mAbs exhibited weaker binding, consistent with the larger antigenic distance between Wuhan and Omicron (**Fig. 3b**). Aligned with the observation with polyclonal mixtures, convergent mAbs were slightly less likely to bind to Delta (78% vs. 91% for donor-specific) and considerably less likely to bind Omicron (44% vs. 73% for donor-specific; **Fig. 3c**). However, convergent mAbs that did bind generally showed similar or stronger affinity than donor-specific mAbs, particularly toward Wuhan S and Delta S, which has relatively few mutations in its S1 subunit (8 compared to Omicron’s 25; **Fig. 3d**).

Finally, to determine how individual serum mAbs contributed to the neutralizing response after vaccination, we performed neutralization assays using VSV-Wuhan, VSV-Delta, and VSV-Omicron (**Fig. 3a, Fig. S9**). Only S^RBD^ mAbs showed detectable neutralization, with 6 of 14 (43%) achieving half-maximal neutralization (IC_50_) of VSV-Wuhan at concentrations below 10 μg/mL. This observation aligns with serology studies from vaccinated non-CF individuals, where RBD-specific antibodies mediate the majority of neutralization activity[61]. Four mAbs neutralized VSV-Delta and two neutralized VSV-Omicron; one mAb, CF2-14, did not neutralize VSV-Wuhan but showed weak neutralization to VSV-Delta. While the sample size is limited, these results suggest that both convergent and donor-specific mAbs can bind S with high affinity, but convergent mAbs may be more susceptible to escape by VOC mutations.

## Discussion

People with CF are at increased risk of infection with viruses including influenza[2, 3, 5, 7, 8, 16, 17]; but lower incidence of infection and relatively mild infection with SARS-CoV-2 compared to the general population[13–15]. The role of the antibody response in immunity against respiratory viruses in pwCF is not understood in detail due to a dearth of high-resolution repertoire studies. We used Ig-Seq[62] to profile the circulating anti-S IgG repertoire in nine pwCF and resolve the serological repertoire at single-clonotype resolution. Our results suggest, broadly, that pwCF mount robust IgG responses to SARS-CoV-2 which resemble those of the general population. In addition, we identify a convergent response in serum comprising 14.6% of S-reactive antibodies on average, and show that this response is associated with lower variant reactivity both overall and on the individual antibody level.

Serum repertoire analysis revealed that the majority of unique clonotypes elicited against S, and the greater part of all circulating antibodies, were specific to RBD (**Fig. 1**). This immunodominance of RBD on S was observed in early B cell sequencing studies in the general population[25, 26]. In addition, a recent serum repertoire study of healthy vaccinees revealed a similar preponderance of RBD-reactive antibodies in circulation[27]. Our single-antibody characterization showed that all mAbs bound S at < 10 nM affinity (**Fig. 3**), though only 6/14 (43%) of RBD-reactive mAbs were neutralizing, which is also in line with B cell level studies in healthy donors[61]. Nevertheless, serum NT_50_ titers to Wuhan pseudovirus were universally strong (**Fig. 1**), suggesting that the high diversity and abundance of RBD-reactive antibodies was sufficient to drive neutralization overall. Clustering all circulating antibodies from pwCF with the hCoV AbDab[57] revealed that convergent S-reactive antibodies between pwCF and healthy donors were observed across all donors (**Fig. 2**). Taken together, these data indicate that pwCF exhibit similar vaccine responses to non-CF individuals both at the repertoire and single-antibody level.

B cell repertoire analysis studies have extensively described convergent (also ‘public’ or ‘shared’) S-reactive antibodies[55, 56, 63], which exhibit varying binding and neutralizing potencies[27, 38, 39]. Our study addresses a key gap in literature on convergent antibodies; namely that direct quantification of these antibodies has been lacking, precluding a direct analysis of the functional consequences of convergence[56]. Ig-Seq enabled us to quantify convergent clonotypes; unexpectedly, they comprised up to 24.8% of all circulating IgG by abundance and were observed across all repertoires (**Fig. 2**). Their abundance correlated inversely with serum neutralizing breadth to both Delta and Omicron, and convergent mAbs showed preferential loss of binding at the single-antibody level relative to donor-specific antibodies. Several examples of convergent antibodies have been shown to lose binding to mutant S proteins; for example, neutralizing antibodies encoded by *IGHV3-53/3-66,* which are overrepresented among our group of convergent antibodies, lose binding to K417 mutants which are observed on Delta and Omicron[38]. Similarly, antibodies encoded by *IGHV1-24* neutralize Wuhan but lose binding in the face of R145 mutations, which are present on Omicron S[34].

Our results are limited by the small number of donors and the lack of a direct non-CF comparator group of vaccinees. Within these limits, the preferential variant escape we observed at both the single-antibody and serum level is consistent with a model in which convergent antibodies concentrate the humoral response on a small number of shared epitopes. Because these antibodies are abundant and arise in many individuals, escape mutations at those epitopes would benefit the virus across a broad swath of the population. Such a model requires convergent antibodies to make up a non-negligible fraction of circulating repertoires, which aligns with our data. Whether this translates into measurable selective pressure on circulating virus will require larger and longitudinal cohorts.

mAbs expressed from convergent clonotypes trended towards stronger Wuhan binding compared to donor-specific mAbs (**Fig. 3**); convergent antibodies that bound Delta did so with higher affinity than donor-specific mAbs. This, along with the low overall SHM observed among convergent antibodies (**Fig. 2**), suggests that convergent responses arise from naïve B cells encoding antibodies with strong germline affinity for S. Recent results support that similar antibodies are observed in the context of HIV gp120[64] and ebolavirus glycoprotein[36]. We suspect that understanding epitopes targeted by the most common convergent antibodies may reveal sites that are more likely to be mutated as the virus continues to circulate. We speculate that the effect of convergent antibodies may be less apparent when recall antibody responses dominate, as in the context of influenza; indeed, convergent responses to influenza have been less frequently reported[65]. As an increasing fraction of the population gains repeated exposure to SARS-CoV-2 through vaccination or infection, the contribution of convergent clonotypes to the serological response beyond first exposure becomes increasingly relevant, since it is not yet clear whether boosting or breakthrough infection raises or diminishes their abundance. Mapping circulating antibody responses to current and future variants may enable the design of booster immunogens that account for mutations at the sites targeted by convergent antibodies, lowering the potential for viral escape, and tracking convergent clonotypes may serve as a tool for anticipating escape mutations or for designing immunogens that steer responses toward less commonly targeted regions.

## Materials and Methods

### Sample Collection

All study protocols were approved by the Institutional Review Boards at Dartmouth College and Dartmouth-Hitchcock Medical Center. Informed consent was obtained from the participants. Blood was taken 7 days and 28-31 days after the second dose of a two-dose mRNA vaccine series with BNT162b2 or mRNA-1273 vaccines. Samples were collected under study number 02000827.

### High-throughput Sequencing of V_H_ and V_L_

500 ng of RNA was extracted from peripheral blood mononuclear cells (PBMC) taken 7 days after second vaccine dose. RNA was reverse transcribed using SuperScript IV enzyme and Oligo(dT) primer (both Invitrogen) according to manufacturer instructions. V_H_ and V_L_-specific transcripts were amplified using the FastStart High Fidelity PCR System (Roche) with gene-specific primers[66]. V_H_ and V_L_ amplicons were sequenced using the Illumina MiSeq platform, with adapters added via two PCR amplifications. Sequences were annotated using MiXCR 2.1.5. IMGT library v3 was used to annotate V_H_ and V_L_ sequences.

### V_H_:V_L_-paired Sequencing

Paired heavy and light chain sequencing from the B cells of each donor’s PBMC samples was done as previously described[67]. Briefly, PBMC were isolated as single cells inside emulsion using a custom flow-focusing device. Droplets contained lysis buffer and poly(dT) conjugated magnetic beads to capture mRNA from individual cells. Magnetic beads were collected and emulsified once more for single-bead RT-PCR, in which V_H_ and V_L_ reverse transcripts were physically linked. The V_H_:V_L_ amplicons were collected from emulsions, amplified using nested PCR, and sequenced using Illumina MiSeq. V_H_ and V_L_ regions were amplified separately for full-length V_H_ and V_L_ sequencing for the recombinant expression of monoclonal antibodies as previously described[41, 52].

### Preparation of Total IgG from Sera and Antigen Enrichment

Each serum sample analyzed in this study was first passed through a 1 mL Protein G agarose (ThermoFisher, 20397) affinity column in gravity mode. Serum flow-through was collected and passed through the column three times. The column was washed with 20 mL of PBS prior to elution with 5 mL of 100 mM glycine-HCl, pH 2.7. The eluate, containing total IgG from serum, was immediately neutralized with 0.75 mL of 1 M Tris-HCl, pH 8.0. Purified IgG was digested into F(ab’)2 with 25 μg of recombinantly-made IdeS protease per 1 mg of IgG for 4 hr on a rotator at 37°C and then incubated with Strep-Tactin agarose (IBA-Lifesciences, 2-1206-025) for 1 hr to remove IdeS protease.

SARS-CoV-2 S (Wu, GenBank_MN908947, residues 14 to 1208 with HexaPro (R682G, R683S, A684A, R685S, K986P, V987P, F817P, A892P, A899P, and A942P mutations))[68] was recombinantly expressed on Expi293F cells (Invitrogen) and purified by HisPur™ Cobalt resin (ThermoFisher, 89966). SARS-CoV-2 RBD (Wu, S residues 319 to 591) conjugated to monomeric Fc was recombinantly expressed in Expi293F cells and purified with Protein G agarose. Recombinant antigen was immobilized on N-hydroxysuccinimide (NHS)-activated agarose resins (ThermoFisher, 26197) by overnight rotation at 4°C. The coupled agarose resins were washed with PBS, and unreacted NHS groups were blocked with 1 M Tris-HCl, pH 7.5 for 30 min at RT. The resins were further washed with PBS and packed into a 0.8 mL centrifuge column (ThermoFisher, 89868). For each sample, F(ab’)2 was incubated with the individual antigen affinity columns for 1 hr on a rotator. Flow-through was collected, and the column was washed with 5 mL of PBS. Antigen-enriched F(ab’)2 was eluted with 1% (v/v) formic acid in 0.5 mL fractions. 30 μL from each elution fraction, neutralized with 1M Tris-HCl, pH 8.0, and pre-flow-through, flow-through, and wash samples were assayed by indirect ELISA with each antigen. ELISA signal in each elution sample was checked using 1:5000 diluted goat anti-human IgG (Fab specific) HRP-conjugated secondary antibodies (Sigma-Aldrich, A0293-1ML). Elution fractions showing a positive ELISA signal were pooled and concentrated under vacuum to a volume of ∼1 μL. 40 μL of DPBS and 2 μL of Tris-HCl, pH 7.5 were added before neutralizing with 1 M NaOH. The neutralized elution samples were concentrated to 50 μL under vacuum.

### MS Sample Preparation

For each enrichment, elution and flow-through samples were denatured with 50 μL of 2,2,2-trifluoroethanol (TFE) and 5 μL of 100 mM dithiothreitol (DTT) at 55°C for 1 hr, and then alkylated by incubation with 3 μL of 550 mM iodoacetamide (Sigma) for 30 min at RT in the dark. Alkylation was quenched with 892 μL of 40 mM Tris-HCl, and protein was digested with trypsin (1:30 (w/w) trypsin/protein) for 16 hrs at 37°C. Formic acid was added to 1% (v/v) to quench the digestion, and the sample volume was reduced to ∼150 μL under vacuum. Peptides were then bound to a Hypersep SpinTip C-18 (Thermo Scientific, 84850), washed three times with 0.1% formic acid, and eluted with a 60% acetonitrile and 0.1% formic acid solution. C18 eluate was concentrated under vacuum centrifugation and resuspended in 50 μL in 5% acetonitrile, 0.1% formic acid.

### LC-MS/MS Analysis

Samples were analyzed by liquid chromatography-tandem mass spectrometry on an Easy-nLC 1200 (ThermoFisher Scientific) coupled to an Orbitrap Fusion Tribrid (Thermo Scientific). Peptides were first loaded onto an Acclaim PepMap RSLC NanoTrap column (Dionex; Thermo Scientific) prior to separation on a 75 μm × 15 cm Acclaim PepMap RSLC C18 column (Dionex; Thermo Scientific) using a 5%-32% (v/v) acetonitrile gradient over 90 mins at 300 nL/min. Eluting peptides were injected directly into the mass spectrometer using an EASY-Spray source (Thermo Scientific). The instrument was operated in data-dependent mode with parent ion scans (MS1) collected at 120,000 resolution. Monoisotopic precursor selection and charge state screening were enabled. Ions with charge ≥ +2 were selected for collision-induced dissociation fragmentation spectrum acquisition (MS2) in the ion trap, with a maximum of 20 MS2 scans per MS1. Dynamic exclusion was active with a 15-s exclusion time for ions selected more than twice in a 30-s window. Each sample was run three times to generate technical replicate datasets.

### MS/MS Data Analysis

Protein sequence databases were constructed using the V_H_ and V_L_ sequences obtained from each donor. V_H_ and V_L_ sequences with ≥2 reads were concatenated to a database of background proteins comprising a consensus human protein database (Ensembl 73, longest sequence/gene) and a list of common protein contaminants (MaxQuant). Spectra were searched against the database using SEQUEST (Proteome Discoverer 2.4; Thermo Scientific). Searches considered fully tryptic peptides only, allowing up to two missed cleavages. A precursor mass tolerance of 5 ppm and fragment mass tolerance of 0.5 Da were used. Modifications of carbamidomethyl cysteine (static), oxidized methionine (dynamic), and N-formylated lysine, serine, and threonine (dynamic) were selected. High-confidence peptide-spectrum matches (PSMs) were filtered at a false discovery rate of < 1% as calculated by Percolator (q-value < 0.01, Proteome Discoverer 2.4).

Iso/Leu sequence variants were collapsed into single peptide groups. For each scan, PSMs were ranked first by posterior error probability (PEP), then q-value, and finally XCorr. Only unambiguous top-ranked PSMs were kept; scans with multiple top-ranked PSMs (equivalent PEP, q-value, and XCorr) were designated ambiguous identifications and removed. The average mass deviation (AMD) for each peptide was calculated as described[69] using data from elution only. Peptides with AMD > 1.7 ppm were removed. Additionally, only peptides identified in ≥ 2 replicate injections for at least one elution sample were kept as high-confidence identifications.

Peptide abundance was calculated from the extracted-ion chromatogram (XIC) peak area, as described[62]. For each peptide, a total XIC area was calculated as the sum of all unique peptide XIC areas of associated precursor ions. The average XIC area across replicate injections was calculated for each sample. For each antigen dataset, the eluate and flow-through abundances were compared and peptides with ≥5-fold higher signal in at least one elution sample were considered to be antigen-specific.

### Peptide-to-Clonotype Index and Mapping

V_H_ sequences were grouped into clonotypes based on hierarchical clustering as previously described[62]. Cluster membership required ≥90% identity across the CDRH3 amino acid sequence as measured by the edit distance. High-confidence peptides identified by MS/MS analysis were mapped to clonotypes, and only peptides uniquely mapping to a single clonotype were considered “informative” and kept. The abundance of each antibody clonotype was calculated by summing the XIC areas of the informative peptides mapping to ≥4 amino acids of the CDRH3 region. SHM levels for individual clonotypes were calculated by averaging the SHM rates of all the V_H_ sequences within each clonotype that contained the detected CDRH3 sequences. A clonotype was classified as “S^RBD^” if it appeared in both pulldowns and if its corresponding PSM and XIC in the S pulldown were no more than 10-fold higher than in the RBD pulldown. Clonotypes with > 10-fold higher PSM or XIC in S than in RBD were designated S^nonRBD^.

### Convergent clonotype Identification from Ig-Seq Data

All unique CDRH3 sequences (1,308 total) detected in the MS data and corresponding to the 929 serum clonotypes were used for clustering, which was done using a modified version of previously published code[29]. In brief, clustering was done with a deterministic approach, beginning with grouping clonotypes by CDRH3 sequence length, *IGHV* gene family, and *IGHJ* gene. Within each group, CDRH3 sequences were iteratively assigned to clusters based on pairwise Hamming distances, with a threshold of 20% difference in amino acid sequence identity (80% similarity cutoff). Initial clusters were further refined by merging clusters if any sequences between them satisfied the similarity threshold. Convergent clonotypes containing only CDRH3’s from a single repertoire were manually removed.

### Curation of Coronavirus Antibody Database

The Coronavirus Antibody Database[57] was downloaded from the Oxford Protein Informatics Group website; the version used (February 18, 2024, total: 12,916 sequences) is current as of November 2024. The database was manually curated to only include human full-length S-reactive antibodies; specifically, nanobodies, antibodies containing non-human genes, and antibodies without available *IGHV* and *IGHJ* gene data were filtered out of the dataset. Antibodies without available full length heavy chain variable sequences were also removed. Any engineered sequences, those recovered from display libraries, and those originating from sources other than B cells taken from SARS-CoV-2 infected or vaccinated individuals were removed. Finally, only sequences known to bind at least one variant of SARS-CoV-2 were kept.

### Circos Plots of Convergent clonotypes

Circos plots were generated in R v4.3.0 using the circlize package[70]. Each sector of the plot represented an individual antibody clonotype which clustered with at least one clonotype from a distinct pwCF serum repertoire. Sector widths were scaled proportionally to the relative abundance of the clonotype in the serum repertoire of the corresponding donor. Clonotypes that clustered together, as described above, were linked manually within the plot to highlight shared features between clonotypes. Donor-specific groups were color-coded according to a predefined palette to visually distinguish contributions from different individuals.

### Recombinant mAb Expression and Purification

Selection of antibody sequences for recombinant expression was based on the combination of V_H_:V_L_-paired databases and proteomics data. First, we identified antibody clonotypes found in the proteomic analysis and searched for the same clonotype in the V_H_:V_L_-paired database. Full-length heavy and light chain sequences were then determined from the paired sequencing database. These genes were purchased as eBlocks (Integrated DNA Technologies) or as gene fragments (Twist Bioscience) and cloned into the pcDNA3.4 vector (Invitrogen). Heavy and light chain plasmids for each monoclonal antibody were transfected into Expi293 cells at a 1:2 ratio. After incubating for 5 days at 37°C with 8% CO_2_, the supernatant containing secreted antibodies was collected by centrifuging cultures at 4000 rpm for 20 min at 4°C. Supernatant was passed over a column containing 1.0 mL Protein G agarose resin three times to ensure efficient capture. After washing the column with 20 mL of PBS, antibodies were eluted with 100 mM glycine-HCl, pH 2.7 and immediately neutralized with 0.75 mL 1 M Tris-HCl, pH 8.0. Antibodies were buffer exchanged into PBS using Pierce™ PES Protein Concentrators with 50 kDa MW cutoff (ThermoFisher 88541).

### ELISA

Half-maximal effective concentration (EC_50_) values and AUC values were used to determine the apparent binding affinities of the recombinant mAbs and sera. SARS-CoV-2 S (‘HexaPro’) and RBD were prepared as described above. SARS-CoV-2 Wuhan NTD was purchased from Immune-Tech (cat. no. IT-002-039p). SARS-CoV-2 Wuhan RBD and S2 domain were obtained from BEI resources (RBD, cat. no. NR-56133; S2, cat. no. NR-53799). B.1.617.2, and B.1.1.529 antigens were purchased from R&D systems (B.1.617.2 S2P, cat. no. 10942-CV; B.1.617.2 RBD, cat. no. 10876-CV; B.1.1.529 S, cat. no. 11060-CV; B.1.1.529 RBD, cat. no. 11056-CV). Costar 96-well ELISA plates (Corning) were coated overnight at 4°C with recombinant antigens, washed with PBST (0.05% Tween-20 in PBS), and blocked with 0.5% BSA in PBS for 2 hr at 4°C. After blocking, serially diluted recombinant antibodies were bound to the plates for 1 hr, followed by 1:500 diluted mouse anti-human IgG (Fc-specific) HRP-conjugated secondary antibodies (ThermoFisher 05-4420) for 1 hr. For serum titering to Wuhan S, the same secondary antibodies were used as for recombinant mAbs. For mAb binding to Wuhan RBD, as well as for total serum titers to Wuhan, B.1.617.2 and B.1.1.529 S antigens, 1:5000 dilution of goat anti-human IgG (Fab-specific) HRP-conjugated secondary antibodies were used instead. For detection, 100 μL 1-Step TMB-ELISA substrate solution (Thermo Scientific, 34028) was added before quenching with 50 μL 1 M H_2_SO_4_. Absorbance was measured at 450 nm using a SpectraMax Paradigm plate reader (Molecular Devices). Data were analyzed and fitted for EC_50_ using a 4-parameter logistic nonlinear regression model in the GraphPad Prism 10 software. CR3022[71] was used as a positive control on Wuhan S/RBD, B.1.617.2 S/RBD, and B.1.1.529 S/RBD plates, and EC_50_ values between plates were normalized using CR3022 EC_50_ values. Influenza 5J8 antibody was used as a negative control on all SARS-CoV-2 antigen binding plates[72].

### Pseudovirus Neutralization Assays

Samples of total serum from vaccinated donors and recombinant mAbs were tested in microneutralization assays using a VSV-SARS-CoV pseudovirus system[73, 74]. Samples were serially diluted 2-fold for total serum (1:12.5-1:3200) and 4-fold for recombinant and control mAbs (4.0 μg/mL to 1.0 ng/mL), and incubated with VSV-SARS-CoV-2 Wuhan S, VSV-SARS-CoV-2 B.1.617.2 S, or VSV-SARS-CoV-2 B.1.1.529 (BA.1) S pseudovirus for 1 hr at 37°C before adding to 293T-hsACE2-expressing cells (Integral Molecular). Plates were incubated at 37°C, 5% CO_2_ for 24 hr, after which luciferase activity was measured in cell lysates using the Bright-Glo system (Promega) with a Bio-Tek II plate reader. Percent neutralization was calculated as 100 - (mean RLU test wells/mean RLU positive control wells) × 100 and used to determine the 50% neutralization titers for serum (NT_50_) and half-maximal inhibitory concentrations for monoclonal antibodies (IC_50_).

### Statistical Analysis

All statistical analyses were performed using GraphPad Prism 10.0 (GraphPad Software, San Diego, CA). All the statistical tests performed are described in the figure legends, and correlations were considered significant at a *p*-value of < 0.05 (\**p* < 0.05, \*\**p* < 0.01, \*\*\**p* < 0.001, ****p < 0.0001, n.s.; not significant).

## Data and Code Availability

The raw proteomic data and high-throughput sequences have been deposited in MassIVE (https://massive.ucsd.edu/ProteoSAFe/static/massive.jsp) under accession ID MSV000100855. Code used for identifying convergent clonotypes in this study is available upon reasonable request. Nucleotide sequences of individual monoclonal antibodies characterized in this paper are deposited at GenBank (accession numbers GenBank: PX977001 - PX977038, and PX977087, PX977088).

## Acknowledgements

The research was supported by Cystic Fibrosis Foundation grant number STANTO19R0. Human subjects were enrolled and samples were collected through the CF Translational Research Core which is funded by BOMBER240 (PI: Bomberger). We are grateful for support from bioMT at Dartmouth through NIH NIGMS Grant P20 GM113132 and Immune Monitoring and Flow Cytometry Resource (IMFCSR) at the Norris Cotton Cancer Center at Dartmouth through NCI Cancer Center Support Grant 5P30 CA023108-41.

---

**Figure S1.**
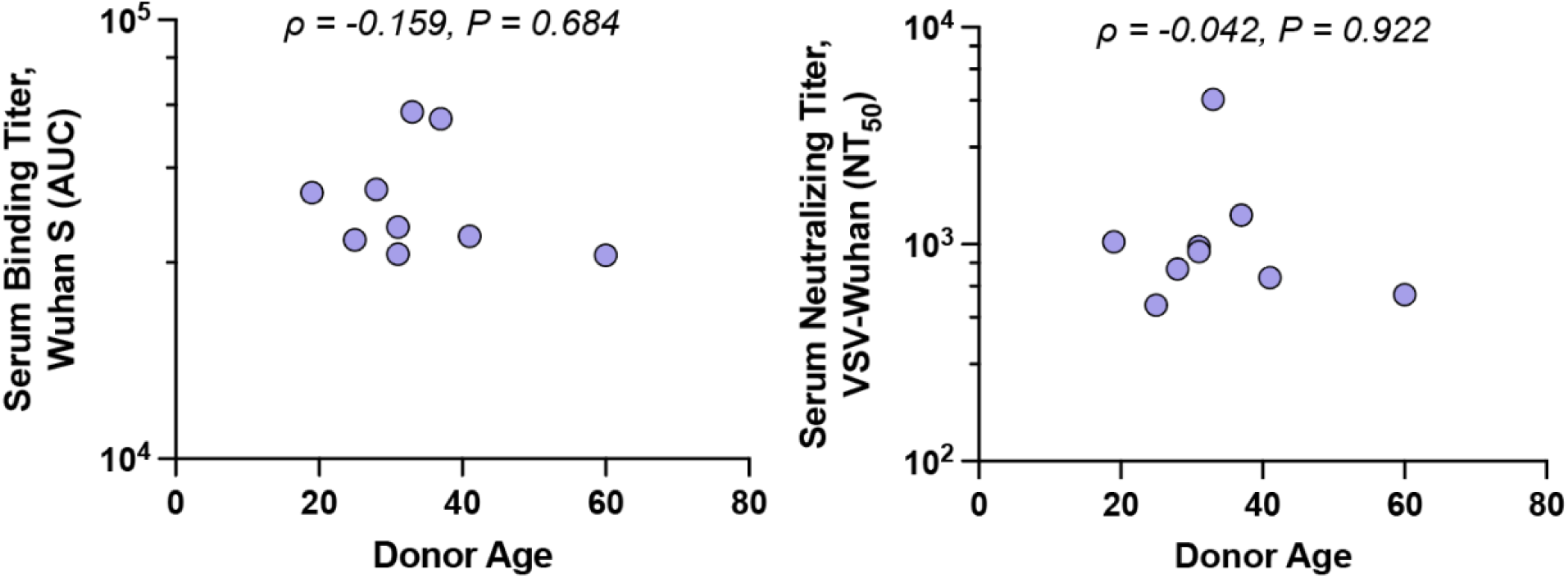
Repertoire features are independent of donor age. **a.** Spearman correlations between donor age and S-reactive serum binding titer. **b.** Spearman correlations between donor age and serum NT_50_ titers to VSV-Wuhan.

**Figure S2.**
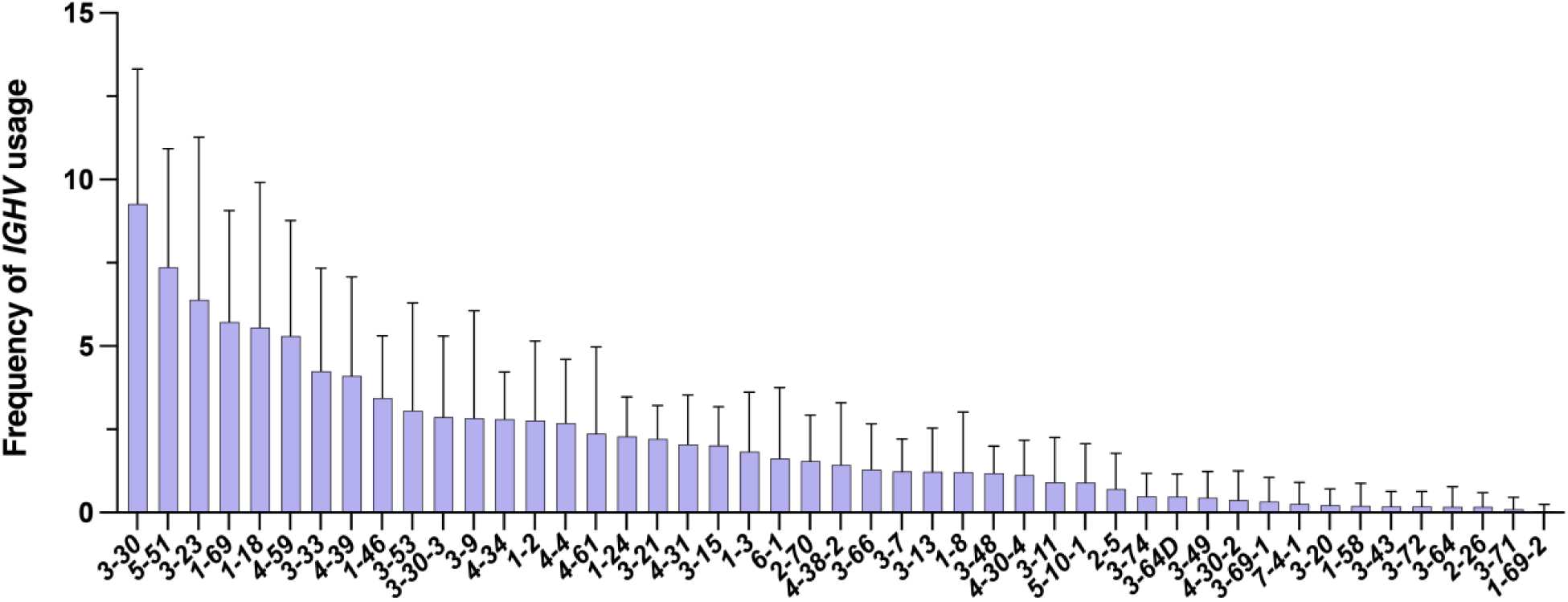
*IGHV* usage among S-reactive serum IgG clonotypes. Gene usage frequency is expressed as percentage of total unique serum clonotypes identified. Bars represent mean frequency across 9 donors, and error bars represent SD.

**Figure S3.**
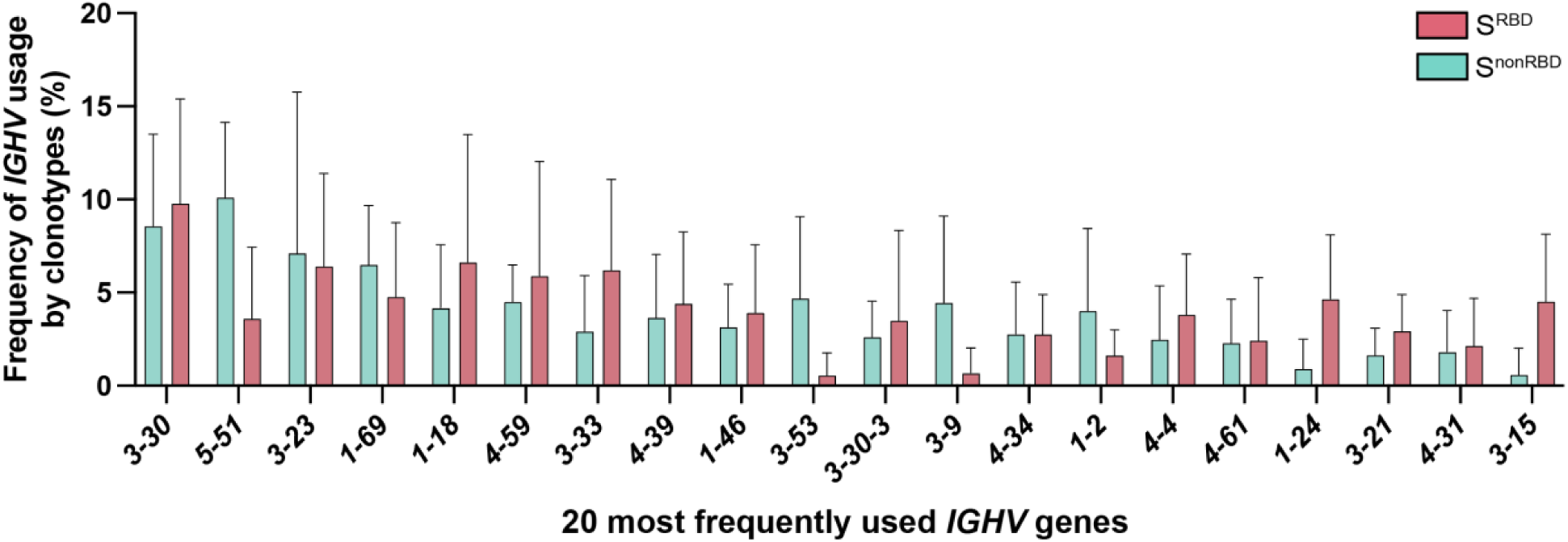
*IGHV* usage among S^RBD^ and S^nonRBD^ clonotypes. Top 20 most frequently used *IGHV* across all clonotypes are shown. Frequencies were calculated for each individual repertoire, and each bar indicates distribution across 9 donors. Error bars indicate SD. No statistically significant differences between S^RBD^ and S^nonRBD^ clonotype frequency were observed among any individual gene. Statistical significance was measured by multiple Wilcoxon tests using Holm-Šidák correction for multiple comparisons, with a *P-*value threshold of 0.05.

**Figure S4.**
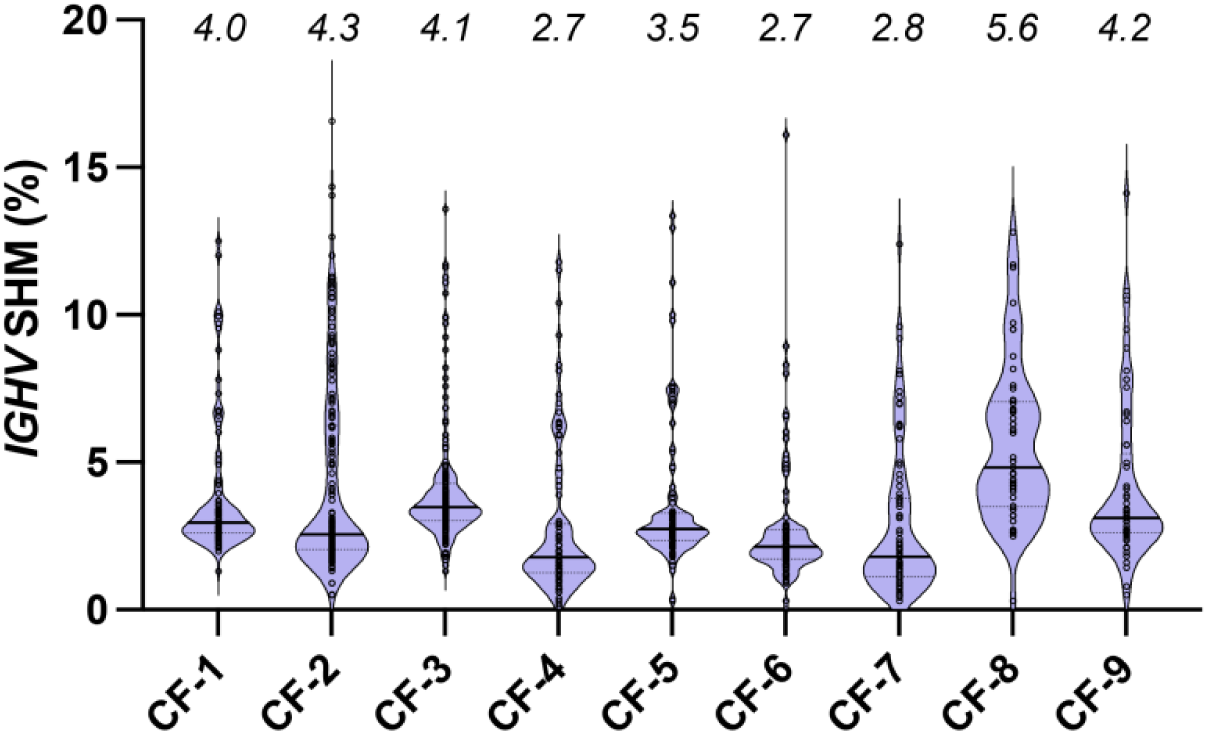
Somatic hypermutation rates of all clonotypes by donor. *IGHV* somatic hypermutation (SHM) levels of serum clonotypes. Numbers above each violin plot represent averages for each donor; solid and dotted lines represent median and quartiles.

**Figure S5.**
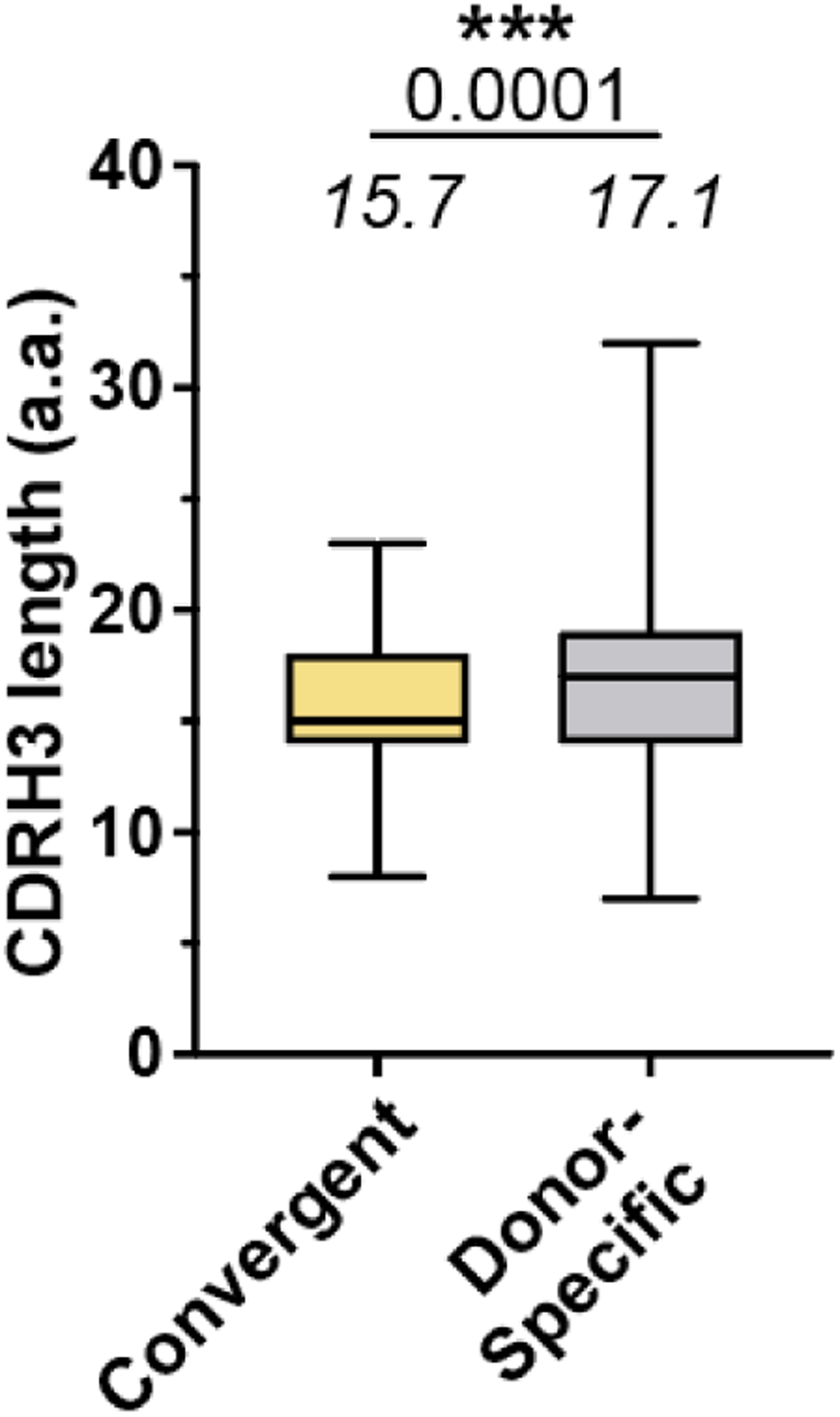
Comparison of CDRH3 amino acid lengths of convergent (yellow) and donor-specific (gray) clonotypes. Box represents first and third quartiles around the median (center line); whiskers range from minimum to maximum. Average values and *P* value are indicated; statistical significance was measured by Mann-Whitney test.

**Figure S6.**
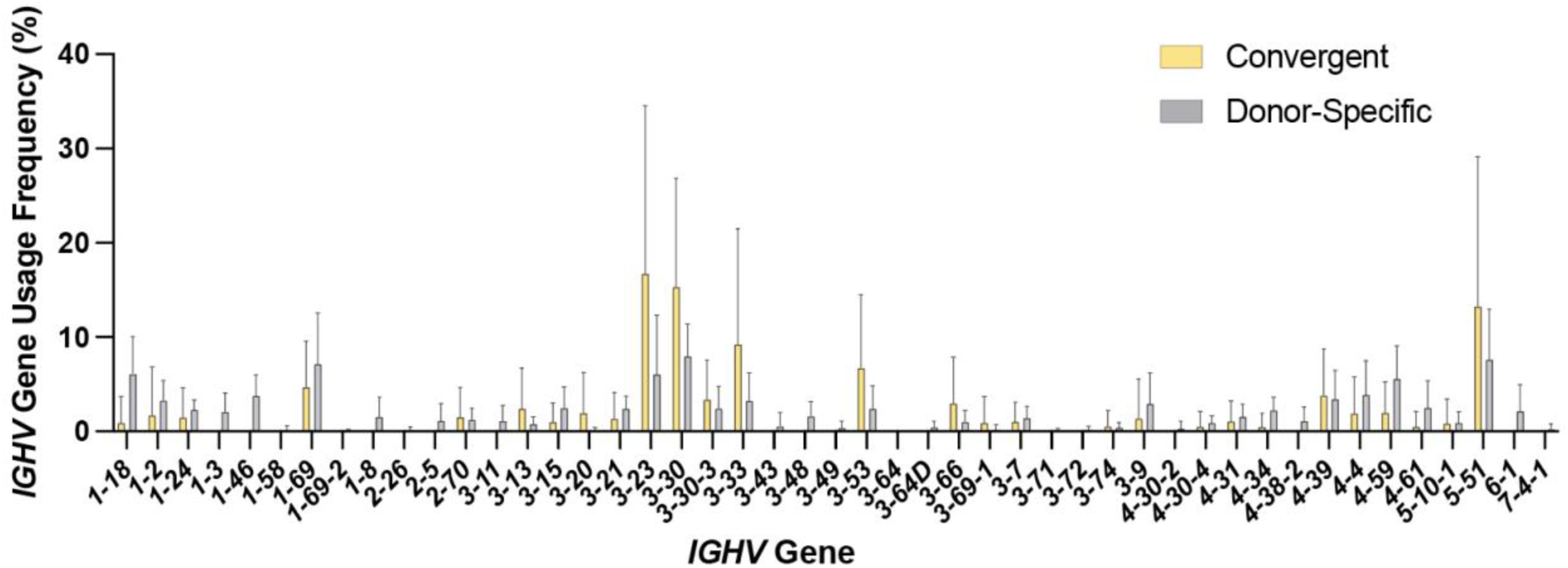
*IGHV* usage by convergent and donor-specific antibodies. Frequency distributions of gene usage by clonotypes classified as convergent (yellow) or donor-specific (gray). Error bars indicate SD.

**Figure S7.**
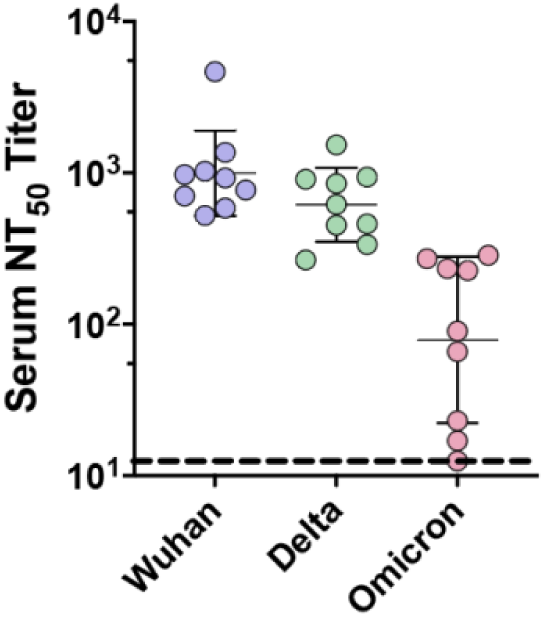
Serum neutralization of VOCs. Total serum half-maximal neutralizing titers (NT_50_) to Wuhan, Delta and Omicron pseudoviruses. Lines and error bars indicate geometric mean and geometric SD. Dashed line at bottom indicates serum dilution of 12.5, the lowest dilution used for titering. One donor with no detectable Omicron NT_50_ titer was assigned an NT_50_ titer of 12.5.

**Figure S8.**
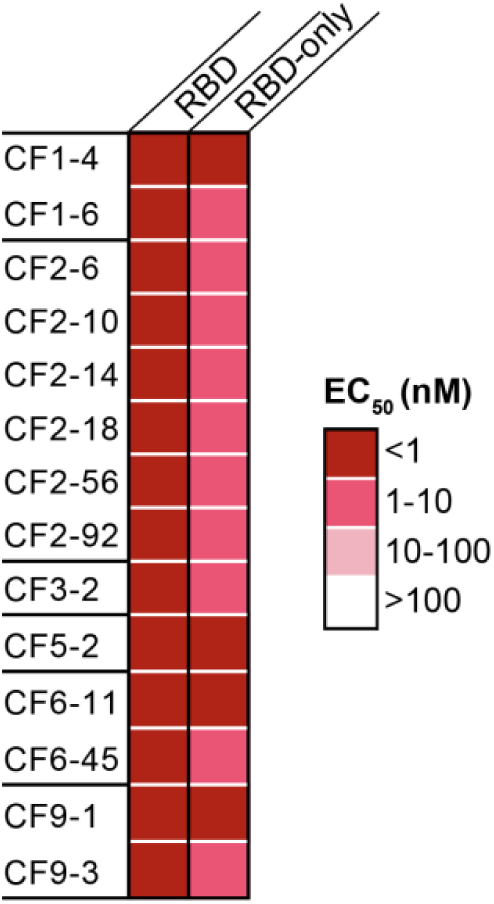
Binding of S^RBD^ mAbs to the RBD-SD1 construct used for Ig-Seq affinity purification (“RBD”), as well as a construct lacking SD1 (“RBD-only”).

**Figure S9.**
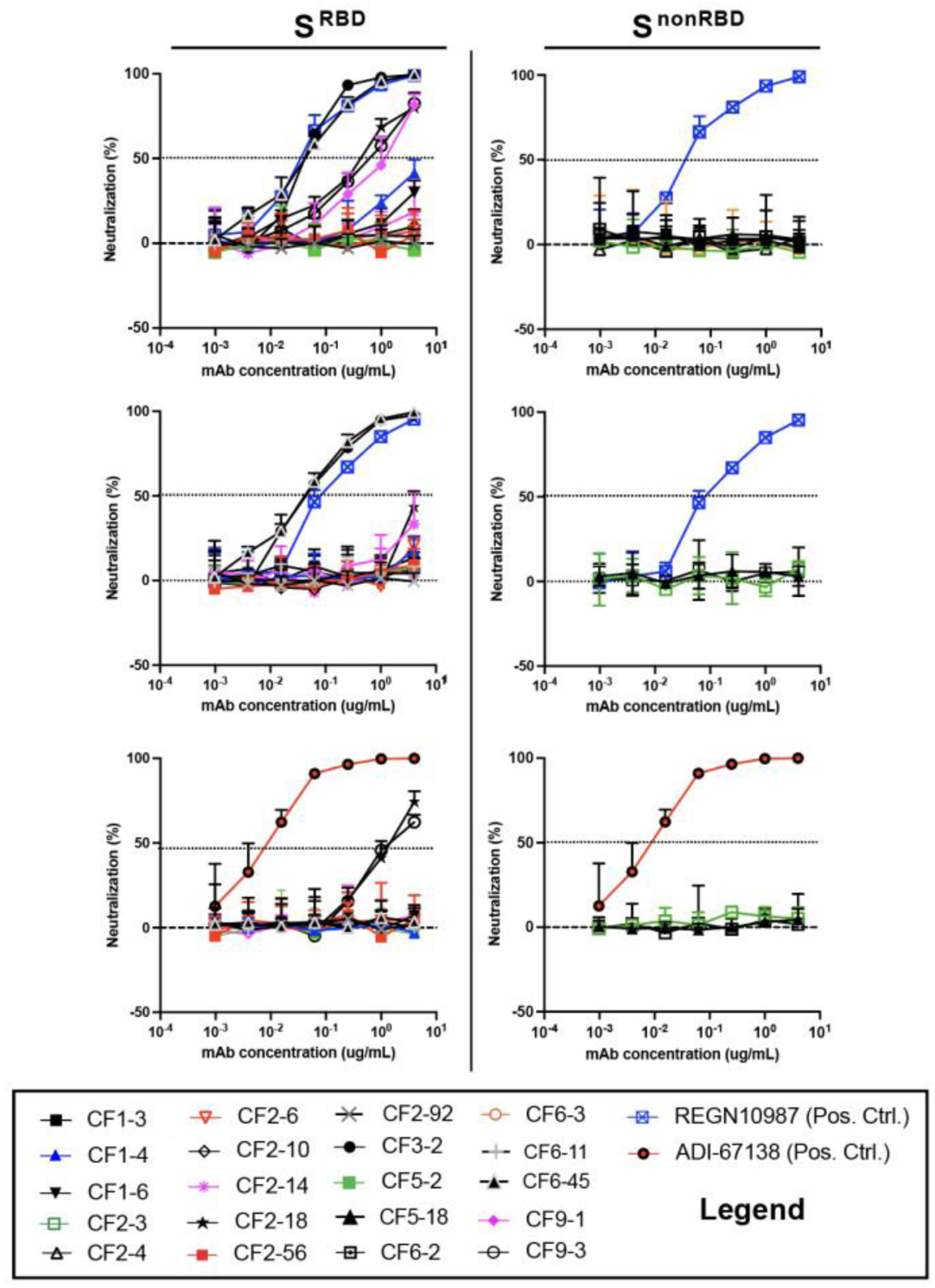
mAb neutralization against Wuhan and VOC pseudoviruses. From top to bottom, neutralization against Wuhan, Delta and Omicron pseudoviruses. S^RBD^ and S^nonRBD^ mAbs are displayed to the left and right of the center line. Dotted lines at 50% are shown to approximate IC_50_ values.

**Table S1.** Vaccination and sample collection information for the pwCF involved in the study.

| Donor ID | Age | Vaccine Received | Dose 1 Date | Dose 2 Date | PBMC Collection Date | Serum Collection Date |
| --- | --- | --- | --- | --- | --- | --- |
| CF-1 | 25 | BNT162b2 | 3/1/21 | 3/24/21 | 3/31/21 | 4/22/21 |
| CF-2 | 41 | BNT162b2 | 3/16/21 | 4/6/21 | 4/13/21 | 5/5/21 |
| CF-3 | 37 | BNT162b2 | 3/15/21 | 4/22/21 | 4/29/21 | 5/20/21 |
| CF-4 | 33 | mRNA-1273 | 3/29/21 | 4/26/21 | 5/3/21 | 5/24/21 |
| CF-5 | 19 | BNT162b2 | 3/13/21 | 4/2/21 | 4/9/21 | 4/30/21 |
| CF-6 | 60 | BNT162b2 | 3/18/21 | 4/15/21 | 4/22/21 | 5/13/21 |
| CF-7 | 31 | mRNA-1273 | 3/24/21 | 4/21/21 | 4/28/21 | 5/19/21 |
| CF-8 | 31 | BNT162b2 | 4/1/21 | 4/22/21 | 4/29/21 | 5/20/21 |
| CF-9 | 28 | mRNA-1273 | 4/6/21 | 5/4/21 | 5/11/21 | 6/2/21 |

**Table S2.** Relative abundance of convergent IgG clonotypes in each pwCF.

| Donor | % Abundance |
| --- | --- |
| CF-1 | 5.9 |
| CF-2 | 10.0 |
| CF-3 | 12.5 |
| CF-4 | 17.8 |
| CF-5 | 3.2 |
| CF-6 | 11.0 |
| CF-7 | 1.6 |
| CF-8 | 2.3 |
| CF-9 | 9.7 |

**Table S3.** Binding of S^nonRBD^ mAbs and SD1-targeting S^RBD^ mAbs to regions of Wuhan S. “RBD-only” indicate RBD construct lacking SD1.

| mAb ID | Convergent? | EC<sub>50</sub> (nM) |  |  |  |  | Specificity |
| --- | --- | --- | --- | --- | --- | --- | --- |
|  |  | S1 | NTD | RBD-only | RBD | S2 |  |
| CF1-3 | No | >100 | >100 | >100 | >100 | >100 | S, unknown |
| CF2-56 | No | 0.53 | >100 | >100 | 0.16 | >100 | SD1 |
| CF5-2 | No | 0.23 | >100 | >100 | 0.15 | >100 | SD1 |
| CF6-2 | No | 1.83 | 2.1 | >100 | >100 | >100 | NTD |
| CF2-3 | Yes | 0.27 | >100 | >100 | >100 | >100 | S1, non-RBD |
| CF2-4 | Yes | 7.95 | 8.07 | >100 | >100 | >100 | NTD |
| CF5-18 | Yes | 0.24 | >100 | >100 | >100 | >100 | S1, non-RBD |
| CF6-3 | Yes | >100 | 3.68 | >100 | >100 | >100 | NTD |

